## Supplemental Figure 1 for "Evaluation of methods incorporating biological function and GWAS summary statistics to accelerate discovery"

**SUPPLEMENTAL FIGURES**

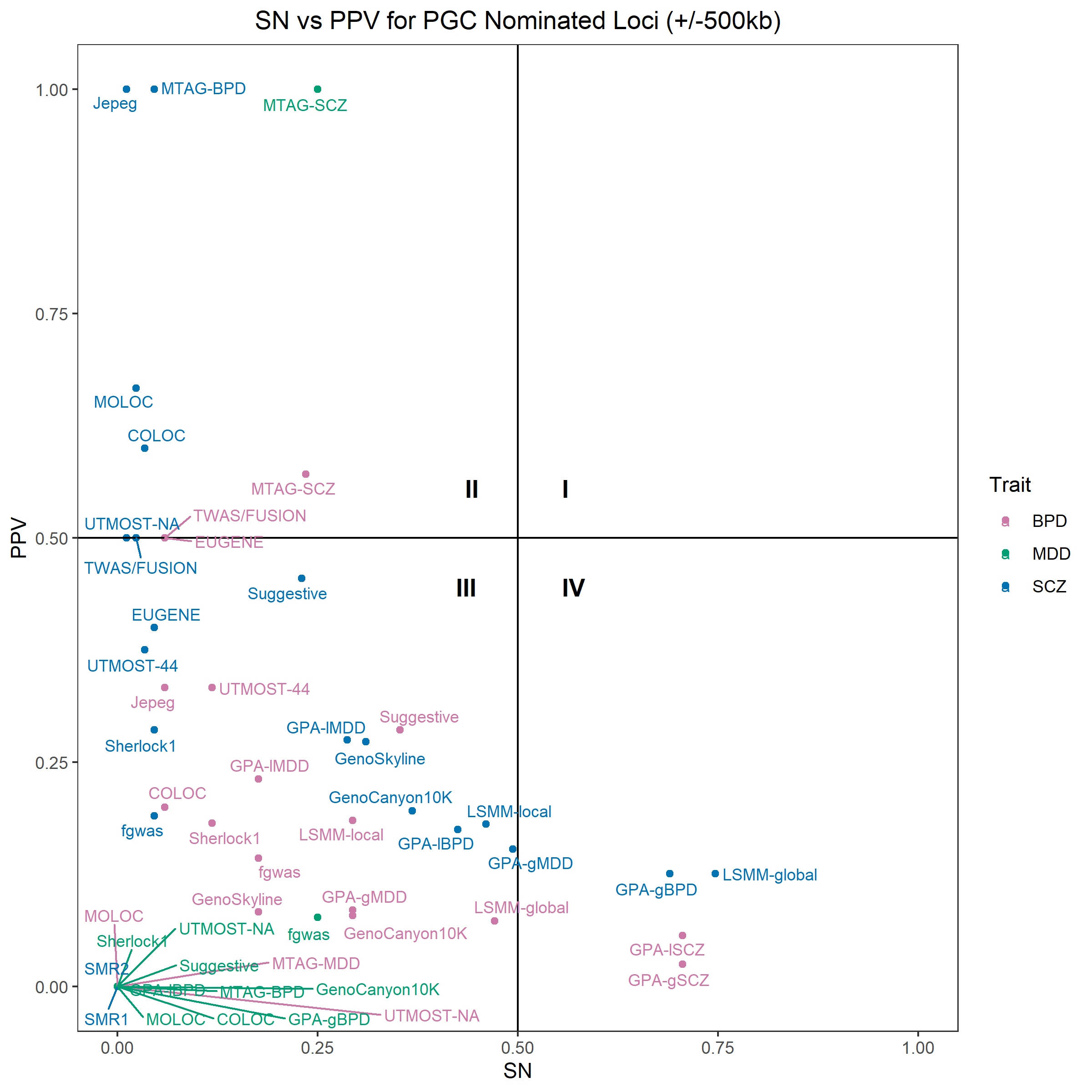

Supplemental Figure 1. Scatterplot of the relationship between sensitivity (SN) and positive predictive value (PPV) for method-psychiatric trait combinations that return nominated variants. SN and PPV were calculated using +/- 500kb overlap criteria and compared to GWAS2 as the gold standard. Horizontal and vertical lines denote SN and PPV of 50%, respectively.
