## Supplemental Figure 2 for "Evaluation of methods incorporating biological function and GWAS summary statistics to accelerate discovery"

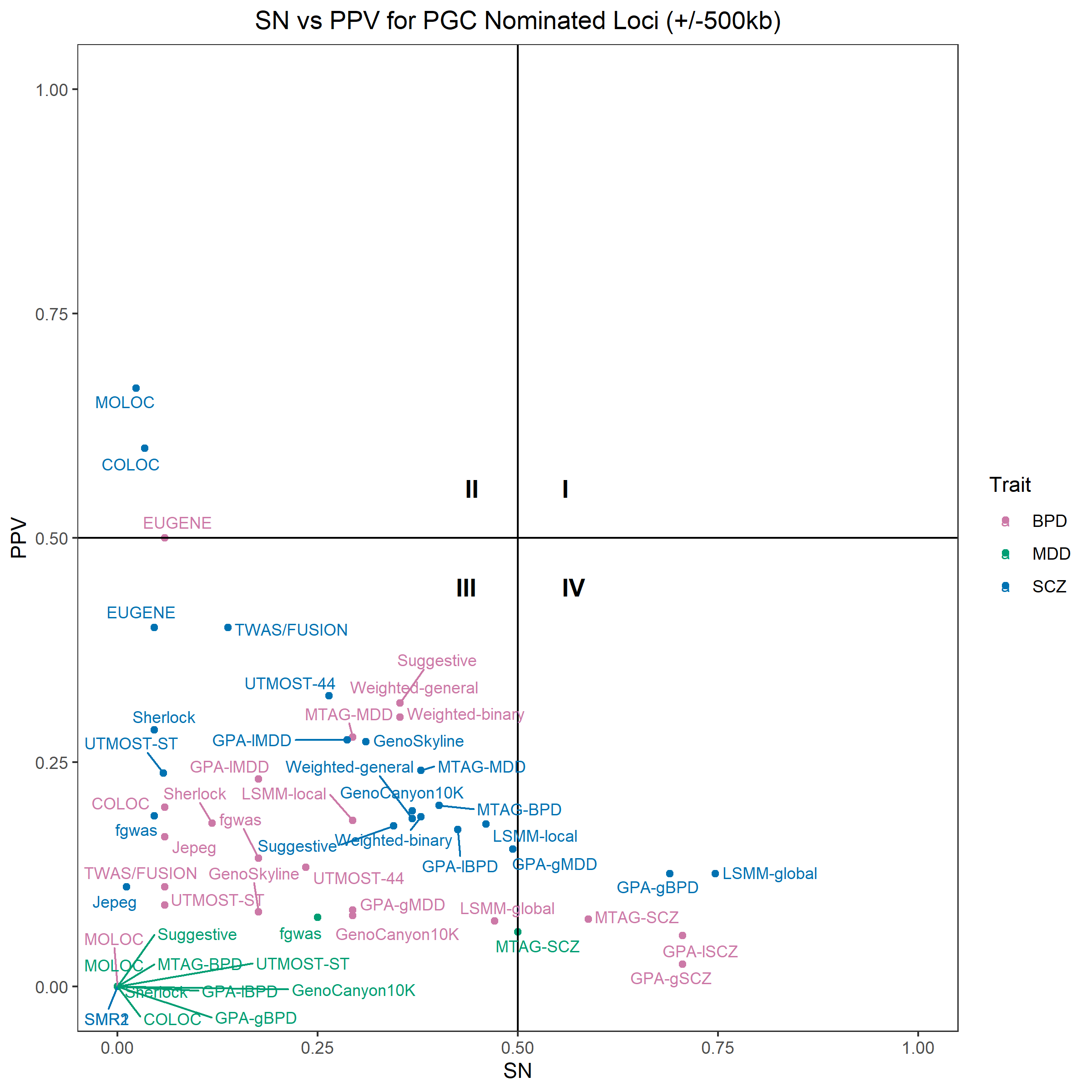

Supplemental Figure 2. Scatterplot of the relationship between sensitivity (SN) and positive predictive value (PPV) for method-psychiatric trait combinations that return nominated variants after using a false discovery rate-based multiple testing correction for the following methods: suggestive, MTAG, Weighted eQTL, JEPEG, TWAS/FUSION, and UTMOST. SN and PPV were calculated using +/- 500kb overlap criteria and compared to GWAS2 as the gold standard. Horizontal and vertical lines denote SN and PPV of 50%, respectively.
