## Supplemental Figure 3 for "Evaluation of methods incorporating biological function and GWAS summary statistics to accelerate discovery"

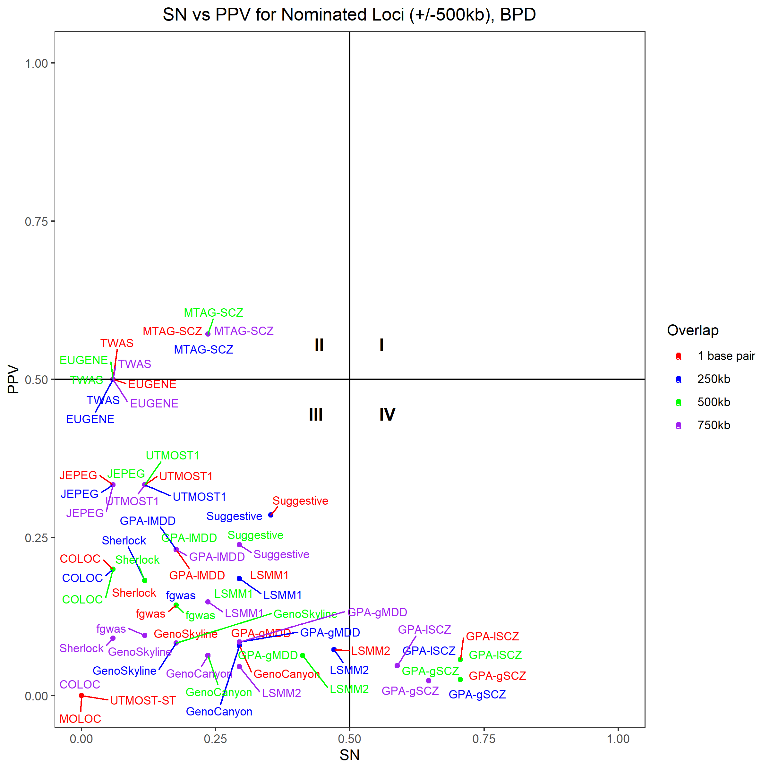

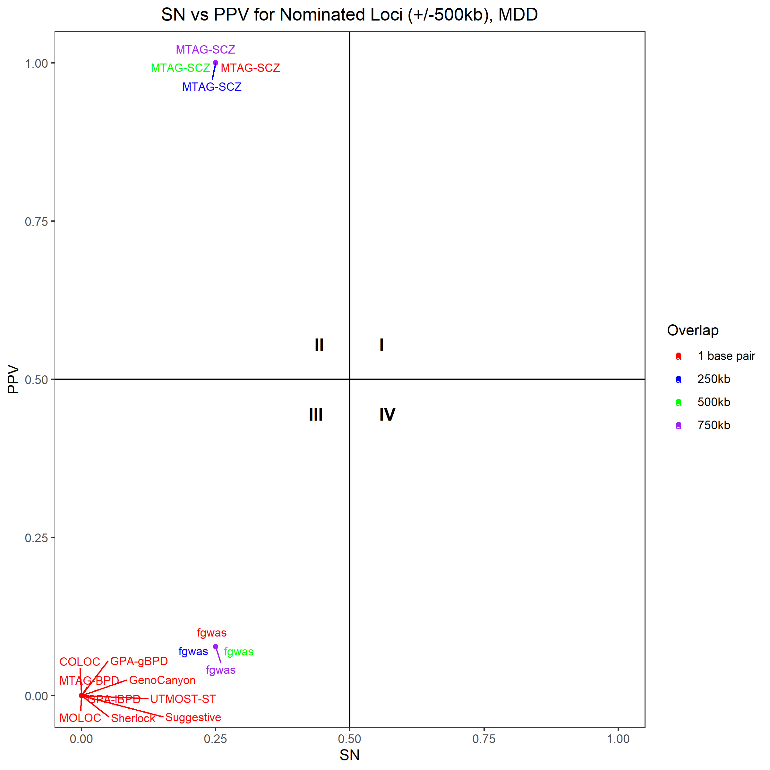

(a) (b)

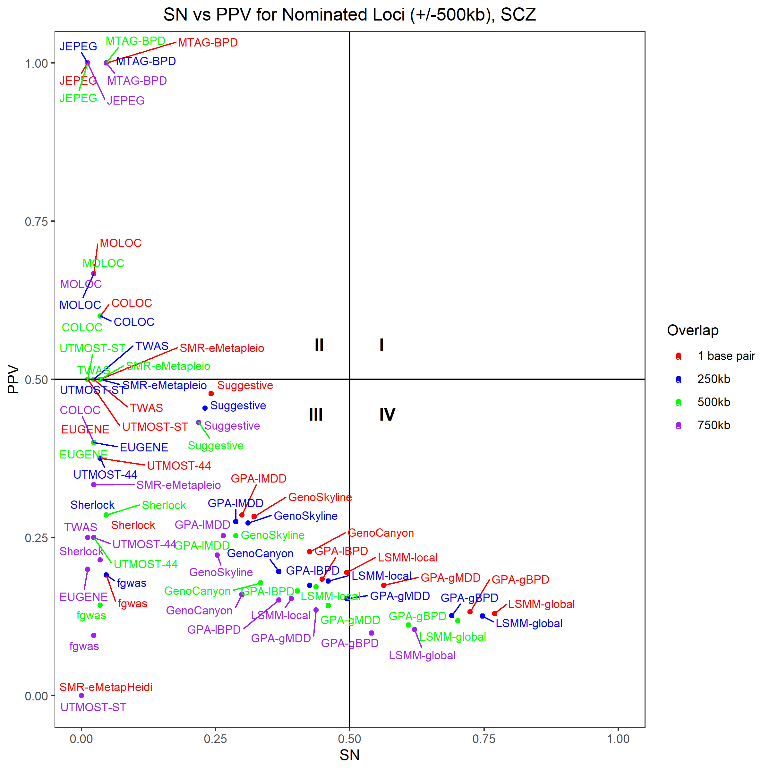

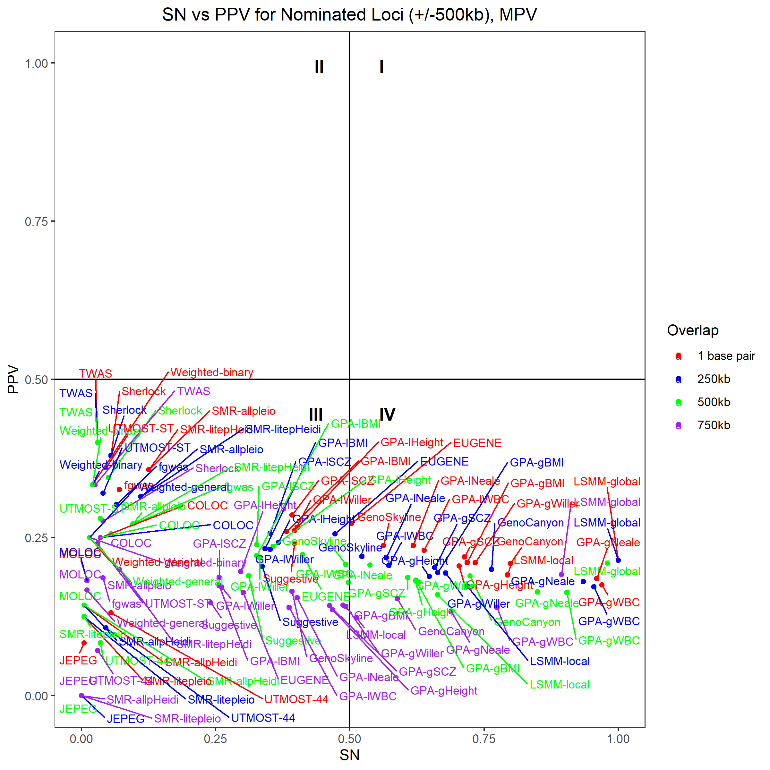

(c)	(d)

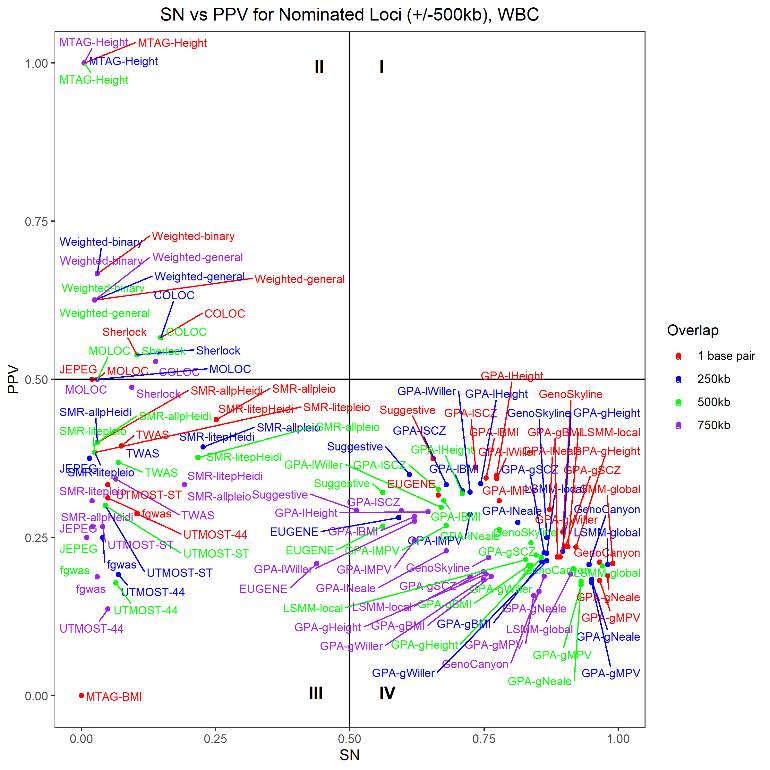

(e)

Supplemental Figure 3. Scatterplots of the relationship between sensitivity (SN) and positive predictive value (PPV) for method-trait combinations that return nominated variants after using different minimum overlap requirements: one base, 250,000 bases, 500,000 bases, and 750,000 bases. SN and PPV were calculated using +/- 500kb loci and compared to GWAS2 as the gold standard for (a) bipolar disorder (BPD), (b) major depressive disorder (MDD), (c) schizophrenia (SCZ), (d) mean platelet volume (MPV), and (e) whole blood cell count (WBC). Horizontal and vertical lines denote SN and PPV of 50%, respectively. For method-trait combinations with SN and PPV of zero at a given minimum overlap requirement, larger overlap requirements are not shown.
