## Supplemental Figure 4 for "Evaluation of methods incorporating biological function and GWAS summary statistics to accelerate discovery"

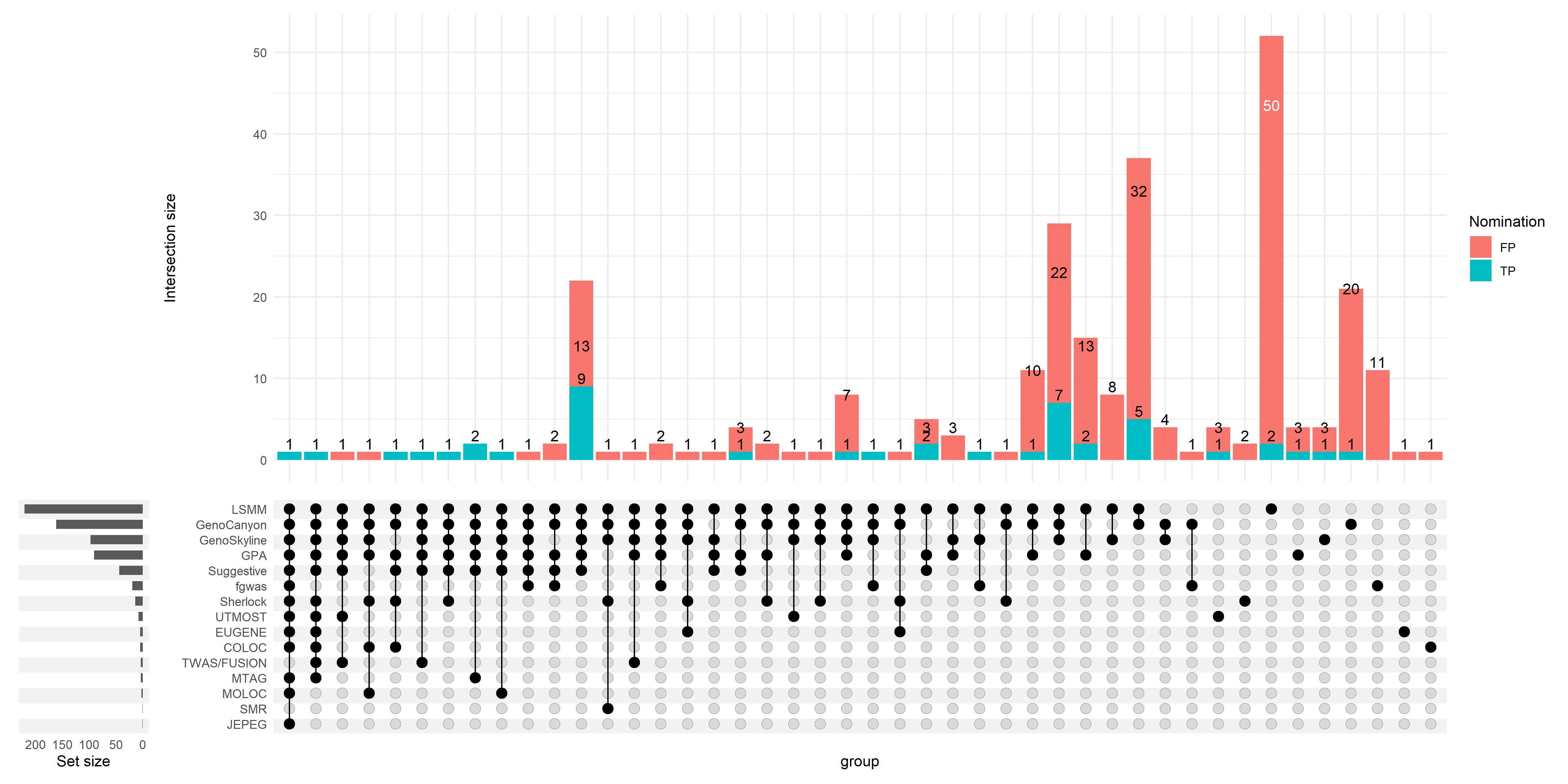

Supplemental Figure 4a. Upset plot of functional weighting methods applied to wave 1 schizophrenia (SCZ1) GWAS.

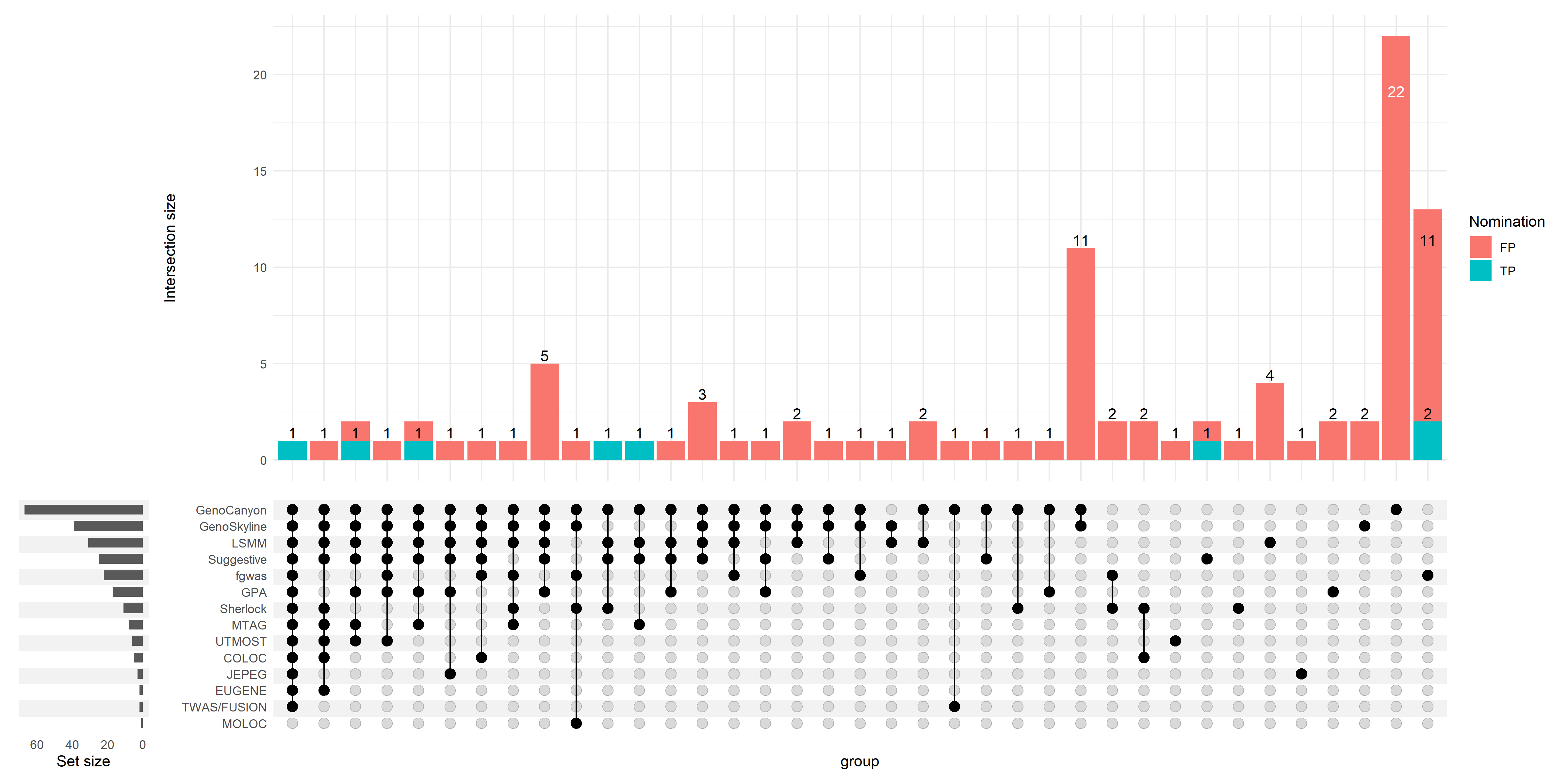
Supplemental Figure 4b. Upset plot of functional weighting methods applied to wave 1 bipolar disorder (BPD1) GWAS.

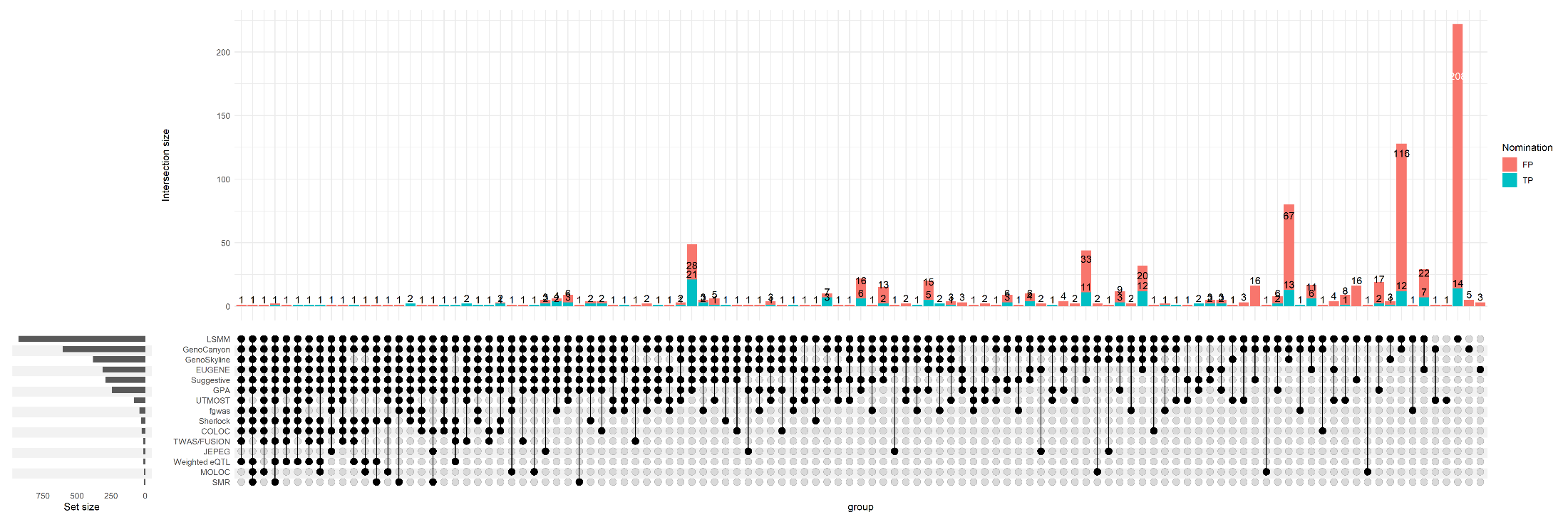

Supplemental Figure 4c. Upset plot of functional weighting methods applied to wave 1 mean platelet volume (MPV1) GWAS.

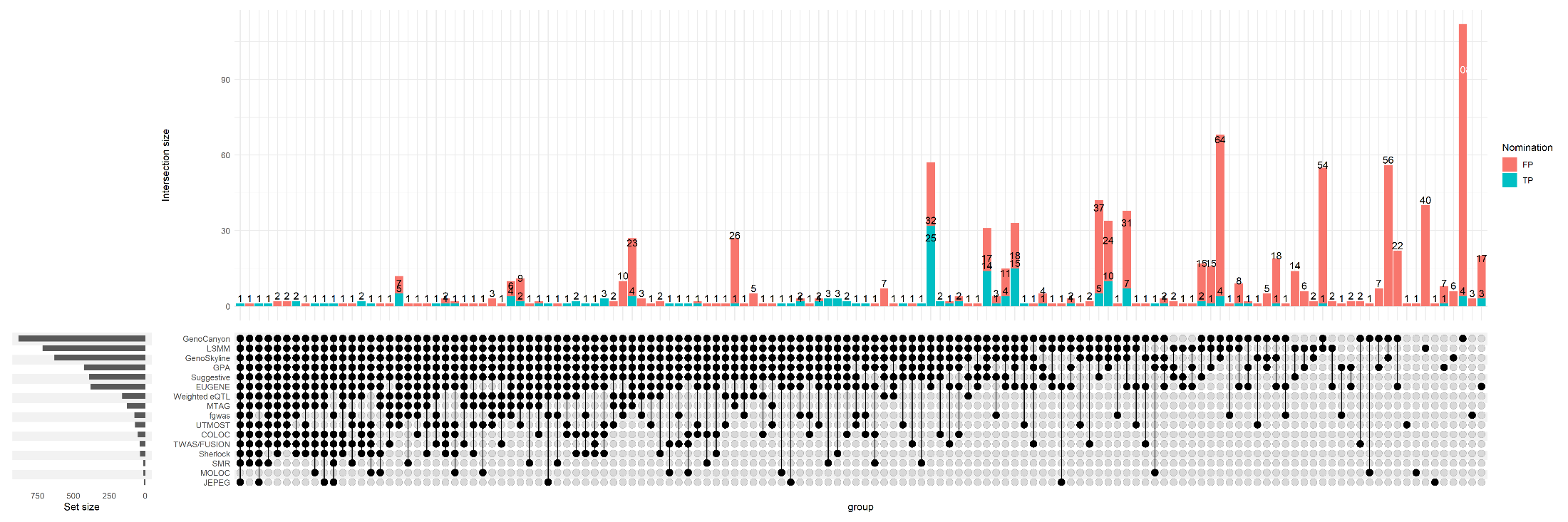

Supplemental Figure 4d. Upset plot of functional weighting methods applied to wave 1 white blood cell count (WBC1) GWAS.

Supplemental Figure 4. UpSet plots of True Positive overlaps among all functional weighting methods applied to SCZ1 (a), BPD1 (b), MPV1 (c), and WBC1 (d), after excluding all loci discovered in the respective GWAS1. Bar plots show the numbers of True Positive and False Positive hits nominated by each method, along with the calculated positive predictive value of each method. To demonstrate the performance of the methods used as directed, SMR was required to have pHeidi < 0.05 and UTMOST was required to be the 44-tissue joint evaluation, except for MDD where the nucleus accumbens single-tissue evaluation was used because the joint evaluation failed to yield any nominations.
